# Molecular Dynamics Analysis of a Flexible Loop at the Binding Interface of the SARS-CoV-2 Spike Protein Receptor-Binding Domain

**DOI:** 10.1101/2021.01.08.425965

**Authors:** Jonathan K. Williams, Baifan Wang, Andrew Sam, Cody L. Hoop, David A. Case, Jean Baum

**Affiliations:** Department of Chemistry and Chemical Biology, Rutgers University, Piscataway, New Jersey 08854; Institute for Quantitative Biomedicine, Rutgers University, Piscataway, New Jersey 08854

**Keywords:** SARS-CoV-2, Molecular Dynamics Simulation, Spike Glycoprotein, Coronavirus, Protein Conformation, Protein Dynamics, Protein Modeling

## Abstract

Since the identification of the SARS-CoV-2 virus as the causative agent of the current COVID-19 pandemic, considerable effort has been spent characterizing the interaction between the Spike protein receptor-binding domain (RBD) and the human angiotensin converting enzyme 2 (ACE2) receptor. This has provided a detailed picture of the end point structure of the RBD-ACE2 binding event, but what remains to be elucidated is the conformation and dynamics of the RBD prior to its interaction with ACE2. In this work we utilize molecular dynamics simulations to probe the flexibility and conformational ensemble of the unbound state of the receptor-binding domain from SARS-CoV-2 and SARS-CoV. We have found that the unbound RBD has a localized region of dynamic flexibility in Loop 3 and that mutations identified during the COVID-19 pandemic in Loop 3 do not affect this flexibility. We use a loop-modeling protocol to generate and simulate novel conformations of the CoV2-RBD Loop 3 region that sample conformational space beyond the ACE2 bound crystal structure. This has allowed for the identification of interesting substates of the unbound RBD that are lower energy than the ACE2-bound conformation, and that block key residues along the ACE2 binding interface. These novel unbound substates may represent new targets for therapeutic design.

## 1. Introduction

The ongoing COVID-19 pandemic is caused by the novel coronavirus SARS-CoV-2 (CoV2), first detected in Wuhan, China in late 2019^1^. The CoV2 genome encodes 29 proteins. Among these proteins is the membrane-anchored spike glycoprotein, a class I membrane fusion protein. The spike protein complex is composed of a homo-trimeric assembly of monomers containing 1273 residues and 22 N-linked glycans^2^ and is responsible for SARS-CoV-2 attachment and entry into host-cells. The virus attachment to human host-cells is mediated by the interaction of the viral Spike protein’s receptor-binding domain (RBD) with the host-cell angiotensin-converting enzyme 2 (ACE2) receptor (**Fig. 1**). Disruption of the binding interface between the Spike protein RBD and host-cell ACE2 receptor would provide a means of preventing SARS-CoV-2 infection at the very first step.

**Figure 1.**
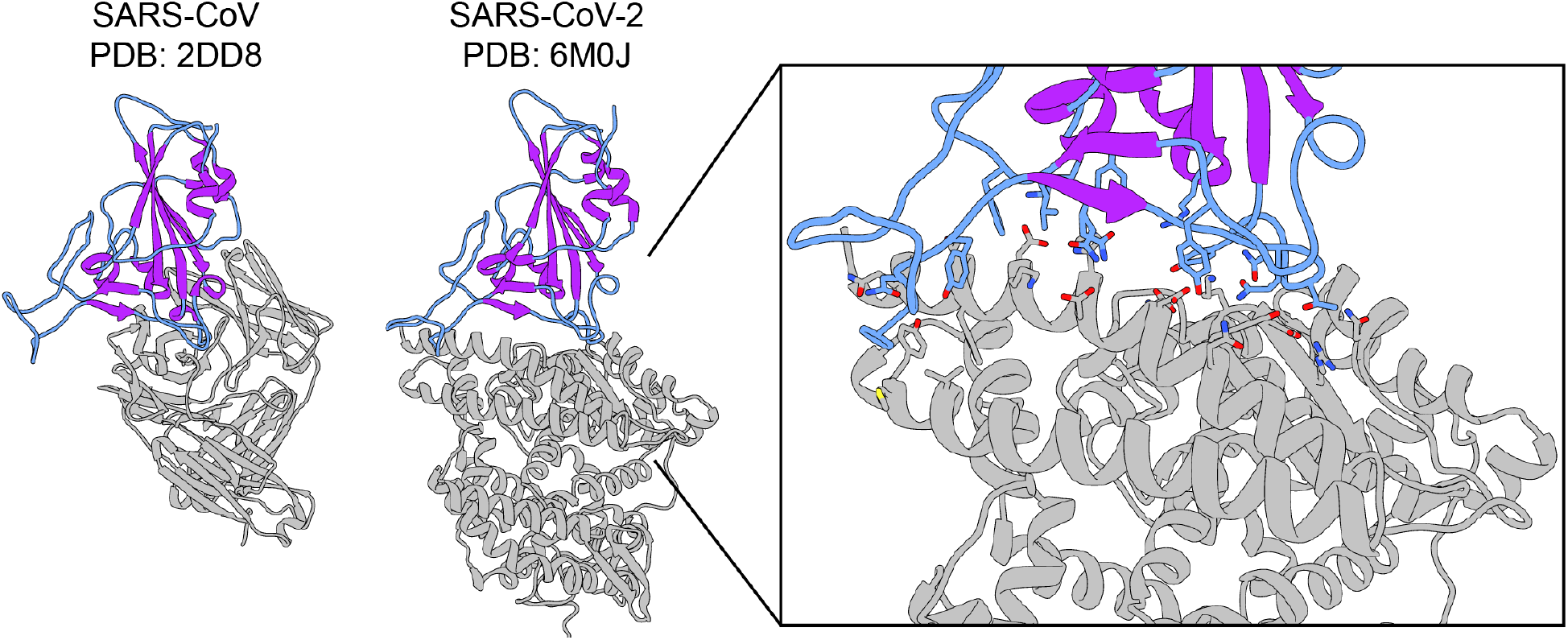
X-ray crystal structures of CoV1 and CoV2 receptor binding domains (RBD) used in MD simulations. X-ray crystal structures of the SARS-CoV RBD in complex with a neutralizing antibody (PDB ID: 2dd8), and the SARS-CoV2 RBD in complex with the ACE2 receptor (PDB ID: 6m0j). The RBDs from these structures are used as starting structures in this work. The RBD is shown in color and the binding partner is in gray. The loop regions are in blue, and the secondary structure elements are in purple, highlighting the large degree of unstructured regions in the RBD. Enlarged inset of the CoV2 RBD-ACE2 binding interface shown on the right. Residue sidechains on the RBD (blue) and ACE2 (gray) that participate in the binding interaction are shown in stick configuration.

Several key aspects of the binding interaction between the Spike protein RBD and the human ACE2 receptor have been characterized for both SARS-CoV-2 and SARS-CoV (CoV1), the coronavirus responsible for a previous pandemic in 2002/2003^3^. These include determining the identity of the binding residues that mediate the interaction between the viral Spike RBD and ACE2, the nature of these residue-level interactions, and the overall strength of the interaction. Both the CoV1-RBD and CoV2-RBD binding sites for ACE2 adopt a similar interface (**Fig. 1**), consisting of long, unstructured stretches of 14 residues which form a range of stabilizing hydrogen bonds, van der Waals contacts, and salt bridges with ACE2^4–6^. The RBD binding interface in general contains 4 loops (**Fig. 2a**) that have the potential to be dynamic and flexible, both in an unbound state as well as after binding to the ACE2 receptor. Several groups have previously investigated the conformational dynamics and flexibility of the RBD when in complex with the ACE2 receptor through molecular dynamics (MD) simulations, identifying that residues 472-490 (*Loop 3*) and residues 495-506 (*Loop 4*) near the binding interface within the receptor binding motif (RBM) as the most flexible regions within the Spike RBD^5,7,8^. In the case of CoV2, three residues within the flexible Loop 3 of the RBD (F486, N487, and Y489) were identified to participate in stabilizing interactions with ACE2^4^. Interestingly, several antibodies developed to target the RBD have also been found to bind to the flexible Loop 3 (**Fig. S1a**). When antibodies interact with Loop 3, the distribution of Loop 3 conformations is greater than when Loop 3 is not part of the binding interface (**Fig. S1b**). This suggests that Loop 3 has an inherent conformational flexibility that is not observed from static structures of the RBD-ACE2 complex. The role that protein dynamics play in mediating protein-protein binding is not only of great importance to understanding the basic mechanisms of binding, but also plays a crucial role in the design of protein binding therapeutics^9,10^.

**Figure 2.**
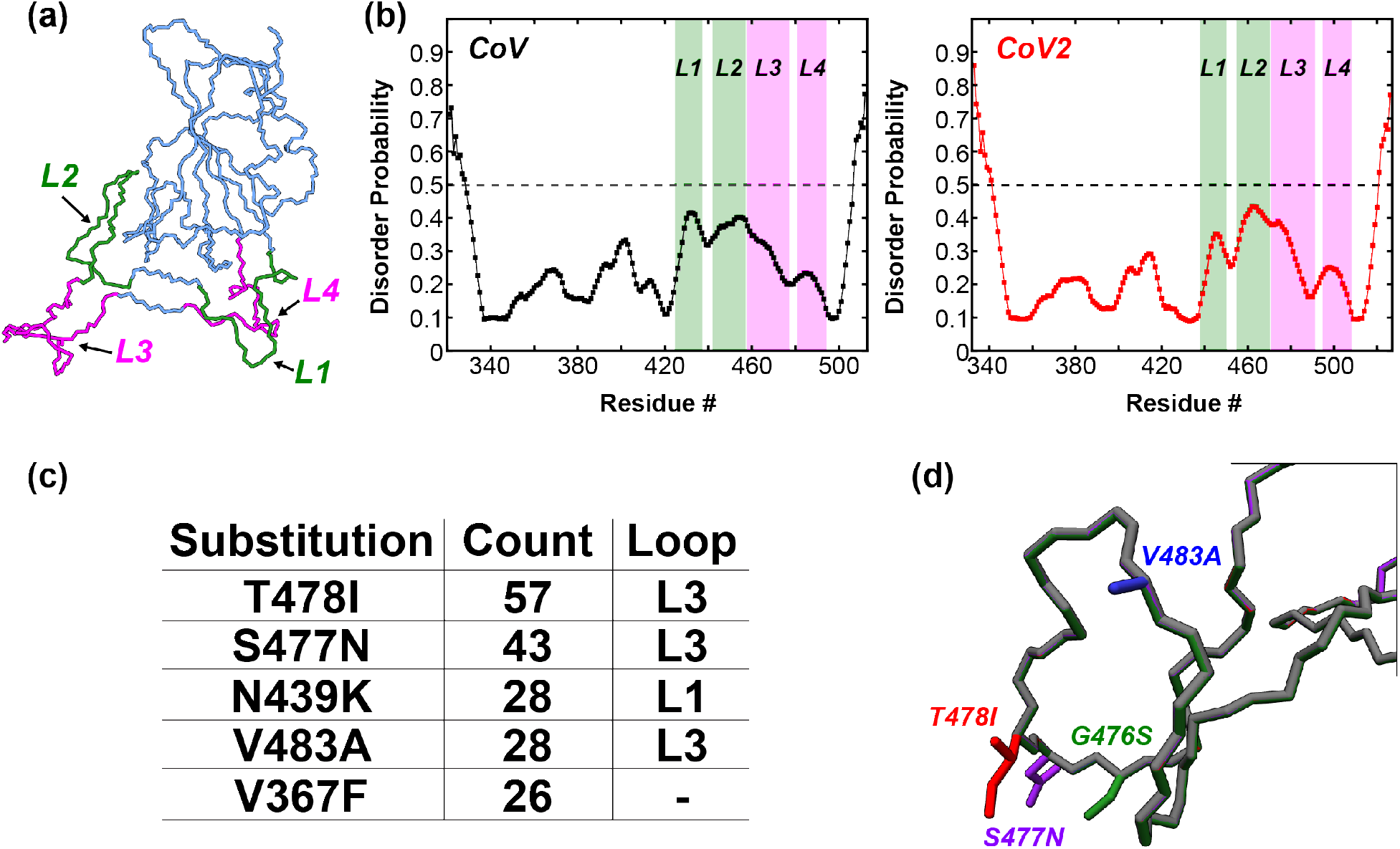
Four loops in the SARS-CoV-2 RBD form the binding interface with ACE2 and harbor several single amino acid substitutions identified during the COVID-19 pandemic. (a) The CoV2-RBD (PDB 6m0j) showing the 4 different loops that make up the binding interface (green and pink). Residues 438-450 (CoV2) or residues 425-437 (CoV) make up Loop 1, residues 455-470 (CoV2) or residues 442-457 (CoV) make up Loop 2, residues 471-491 (CoV2) or residues 458-477 (CoV) make up Loop 3, and residues 495-508 (CoV2) or residues 481-494 (CoV) make up Loop 4. (b) Prediction of natively disordered regions using the Protein Disorder prediction System (PrDOS) webserver^11^ for CoV-RBD (left) and CoV2-RBD (right). PrDOS was used without template-based prediction and thus reports only on the disorder probability of the local amino acid composition. A prediction false positive rate of 5% was used, and values above the 50% threshold (dotted line) indicate regions of predicted disorder. (c) The 5 most common mutations of the RBD identified during the first 6 months of the COVID-19 pandemic, with 4 of these being located in the loop regions of the binding interface. A complete list of RBD mutants can be found in Table S1. (d) Depictions of the side-chains of the mutant residues contained in Loop 3 that are studied in the current work.

Although much progress has been made in understanding the interaction between the Spike RBD and ACE2, what remains to be elucidated is the flexibility and conformational dynamics of the RBD in an unbound state. The internal motions of proteins play a key role in their interactions and functionality, a fact that is often lost in static structures derived from electron microscopy and X-ray crystallography. Understanding the conformational ensemble of RBD states without a binding partner may reveal novel targets not observed in static structures of the RBDs, which will aid in the design of therapeutics targeting this important binding domain. In this work we utilize molecular dynamics simulations to probe the flexibility and conformational sampling of the SARS-CoV and SARS-CoV-2 receptor-binding domains in an unbound state. We focus on the Loop 3 region of the RBD, which contains several residues that participate in stabilizing interactions with ACE2 and is a hot-spot of several common single amino acid mutations that have been identified during the ongoing COVID-19 pandemic. We find that Loop 3 represents a localized area of dynamic flexibility in an unbound state, and our simulations suggest that this flexibility is resilient to perturbation by mutations. Finally, using loop-modeling to probe novel conformations of the Loop 3 region, we have identified interesting substates of the unbound RBD that block the binding interface and are lower energy than the conformation of the RBD bound to ACE2, and thus may represent enticing targets for therapeutic intervention.

## 2. Results

### Microsecond Timescale MD Simulations of Wild-Type CoV1 and CoV2 RBDs Reveal Localized Flexibility in Loop 3

Multi-microsecond molecular dynamics simulations were recorded in order to explore the conformational flexibility and dynamics of the wild-type spike protein RBDs from CoV1 and CoV2. The initial coordinates used in the MD trajectories were taken from the high-resolution structures determined by X-ray diffraction of CoV1-RBD in complex with a neutralizing antibody (PDB: 2dd8) and of CoV2-RBD in complex with the human ACE2 receptor (PDB: 6m0j) (**Fig. 1**). While it contains a well ordered b-sheet core, much of the RBD is unstructured (**Fig. 1**) and in particular 4 different loops make up the binding interface (**Fig. 2a,** *green and pink*) of the RBD with ACE2. Residues 438-450 (CoV2) or residues 425-437 (CoV1) make up Loop 1, residues 455-470 (CoV2) or residues 442-457 (CoV1) make up Loop 2, residues 471-491 (CoV2) or residues 458-477 (CoV1) make up Loop 3, and residues 495-508 (CoV2) or residues 481-494 (CoV1) make up Loop 4. An assessment of the residue level propensity for disorder, using the Protein Disorder prediction System (PrDOS) webserver^11^, indicates that while none of these regions is considered intrinsically disordered, the loop regions of both CoV1-RBD and CoV2-RBD do show an increased disordered propensity (**Fig. 2b**) relative to the rest of the RBD. PrDOS was used without template-based prediction and thus reports only on the disorder probability of the local amino acid composition. A prediction false positive rate of 5% was used, and values above the 50% threshold (dotted line) indicate regions of predicted disorder.

As observed from an analysis of root-mean-square deviation (RMSD) with respect to the starting structure from 4 μs MD trajectories (**Fig. S2a**) both CoV1-RBD and CoV2-RBD remain in a stable equilibrium conformation over the time-course of the MD trajectories, with average RMSD values of 1.42 Å for CoV1 and 1.39 Å for CoV2. However, the RMSD of CoV1-RBD shows several large fluctuations, suggesting that CoV1-RBD is more conformationally flexible than CoV2-RBD. Indeed, this is observed in the calculated per residue root-mean-square-fluctuation (RMSF) profiles (**Fig. 3a**) and in snapshots along the MD trajectory (**Fig. 3b**). The RMSF profiles indicate that the CoV1-RBD overall is more flexible than CoV2-RBD. However, the CoV2-RBD does show a localized area of increased flexibility in residues 369-373 relative to CoV1-RBD. Both CoV1 and CoV2 RBDs have substantial flexibility in the Loop 3 region from ~471-491 (**Fig. 3a, inset**), which is part of the large ACE2 binding interface. The average conformations obtained from the MD simulations of CoV1-RBD and CoV2-RBD are quite similar (**Fig. 3c**), with the major differences localized to Loop 3 centralized around the conserved disulfide bond between residues 480 and 488 (**Fig. 3c, enlargement**).

**Figure 3.**
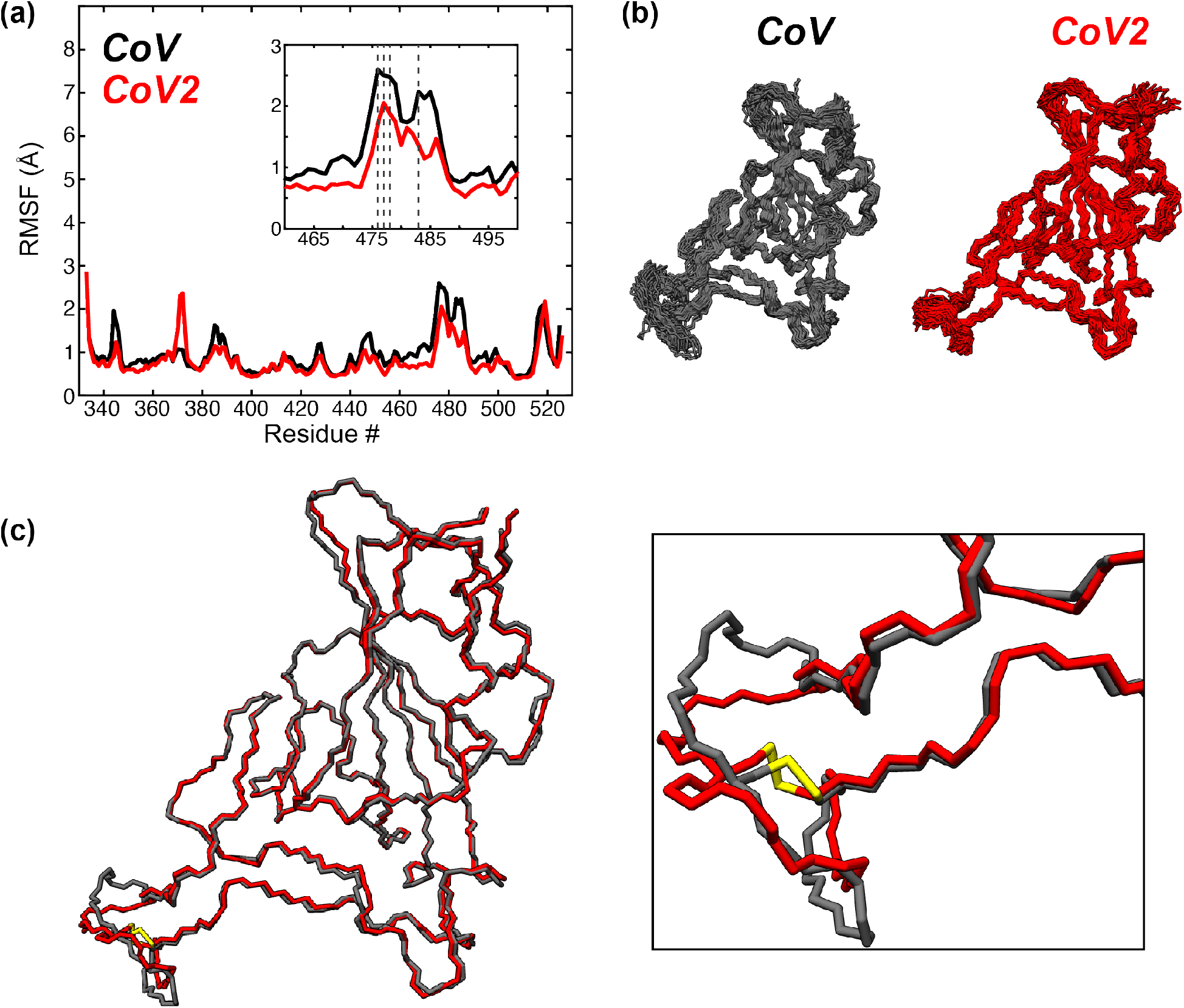
Microsecond timescale MD simulations of wild-type SARS-CoV and SARS-CoV2 RBD. (a) Per residue root-mean-square fluctuation (RMSF) of all backbone (N, CA, C) atoms of SARS-CoV (black) and SARS-CoV2 (red) RBDs from 4 μs MD trajectories. The sequences were aligned, and the CoV2 residue numbering is used as reference for the x-axis. The small inset shows the large fluctuation of the Loop 2 and Loop 3 regions near the binding interface with ACE2, where several mutations in the RBD are clustered (dotted lines). (b) Conformational snapshots throughout the 4 μs MD trajectories. (c) Average conformations of the SARS-CoV (black) and SARS-CoV2 (red) RBDs, with a focus on the disulfide (yellow)-containing loop region that shows large fluctuations over the 4 μs simulation. The colors of the data and models are kept consistent throughout the figure.

### Conformational Flexibility of Loop 3 in the Free RBD

In order to understand the interaction mechanisms of the RBD with binding partners and for design of therapeutics, one needs to understand the conformations accessible in the free state of the RBD prior to binding. The MD simulations starting from the crystal structures show that while Loop 3 does display the highest flexibility within the RBD, on average there is not a large deviation from the starting structure. This might result from being trapped near the starting point of the crystallized ACE2-bound state of the RBD. To better probe the conformational flexibility of the RBD in the free state, we conducted MD simulations of several unique loop-model structures of the Loop 3 region (**Fig. 4**). The KinematicMover algorithm within pyRosetta was used to generate 100 new conformations of the Loop 3 region of the CoV2 RBD (PDB: 6m0j) and 100 new conformations of the Loop 3 region (residues 458-477) of the CoV1 RBD (PDB: 2dd8), making sure to maintain the disulfide bonds (CoV2: C480-C488, CoV1: C467-C474). Five of these new conformations were then chosen at random, and subjected to energy minimization and relaxation protocols in pyRosetta as described in the Methods section. The 5 energy minimized loop-models were used as starting structures for 750 ns of MD simulation, and snapshots from these simulations are shown along the outer edges of **Fig. 4**, along with the average structures from each of the 5 simulations overlaid in the center of the figure and an enlargement of the Loop 3 region shown in the boxes at the bottom.

**Figure 4.**
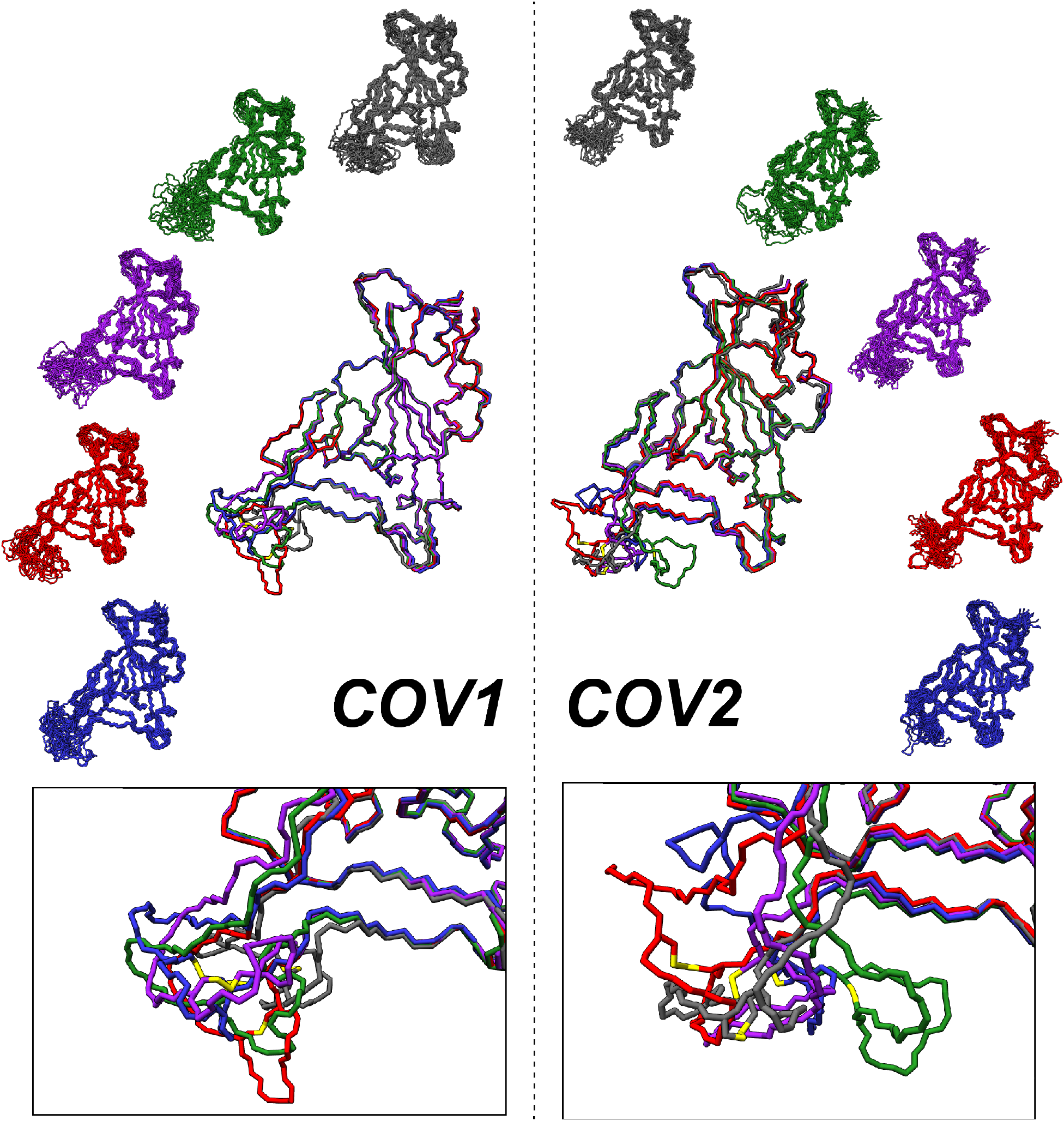
Diverse conformational sampling of Loop 3 from different loop models of the RBD. (Left Side) Snapshots from 750 ns of MD simulations of 5 different loop model structures of the SARS-CoV RBD are shown on the periphery, with the average structure from each simulation overlaid in the center. The bottom box shows the sampling of Loop 3 conformational space from overlaid average structures. (Right Side) Snapshots from 750 ns of MD simulations of 5 different loop model structures of the SARS-CoV-2 RBD are shown on the periphery, with the average structure from each simulation overlaid in the center. The bottom box shows the sampling of Loop 3 conformational space from the overlaid average structures. The different colors of the models in both cases are used to differentiate between the different starting structures used for each simulation.

The loop modeling shows that Loop 3 can take on a vast range of conformations. In general the loop models represent an increase in free energy, but both CoV1-RBD and CoV2-RBD have one loop model that represents an average conformation with lower free energy compared to the crystal structure. The average difference in free energy between the conformations sampled in the simulations from crystal structures versus the simulations of the different loop-models is summarized in **Table 1**. These DG values were calculated as the difference between the average Molecular Mechanics/Generalized Born Surface Area (MM/GBSA) free energy for each loop model simulation and the average MM/GBSA free energy from the 4 μs simulations described in the previous section. The RMSDs of the conformations sampled during the MD simulation with respect to the starting structure show large variation between different loop models for both CoV2 (**Fig. S2c**) and CoV1 (**Fig. S2d**), although the CoV1 loop models (1.8–2.7 Å) are clustered at a smaller RMSD average than the CoV2 loop models (2.0–4.8 Å). To better deconvolute the contribution of Loop 3, the backbone RMSD with respect to the starting structure was recalculated ignoring the Loop 3 residues (**Fig. S2c,d**) and also by only considering the Loop 3 residues (**Fig. S2c,d**). When viewed in this fashion, it becomes clear that the increase in RMSD observed in the loop-modeled simulations relative to the crystal structure simulations is a result of the increased flexibility of the new loop-conformations. Indeed when viewed on a per-residue basis (**Fig. S3**), the overall RMSF profiles of the loop-models maintains the same topology as the wild-type simulations, while displaying drastic increases in the RMSF values of Loop 3 for both CoV1 and CoV2 (maximum RMSF ~8-9 Å, compare to Fig. 3a). The range of conformations probed in the loop modeling, some of which were more energetically favorable than the crystal structure, suggests that Loop 3 is capable of sampling a variety of conformations in solution. Future experimental studies will be necessary to further probe and define the conformational dynamics of the RBD, and especially the Loop 3 region.

**Table 1.** Calculated average Molecular Mechanics/Generalized Born Surface Area (MM/GBSA) changes in free energy compared to the starting structure (see Methods).

| Loop<br>Model | CoV1-RBD | CoV2-RBD |
| --- | --- | --- |
| | $\Delta G$ (kcal/mol) | $\Delta G$ (kcal/mol) |
| 1 (black) | 13.2 | 6.7 |
| 2 (green) | 12.1 | 15.4 |
| 3 (purple) | 31.7 | 14.2 |
| 4 (red) | -0.04 | 6.5 |
| 5 (blue) | 10.4 | -3.5 |

### Analysis of Spike Protein Mutations Accumulated During the COVID-19 Pandemic Highlight a Mutational Hotspot in Loop 3

It is important to consider how mutations perturb the conformation and dynamics of the RBD, especially during an ongoing pandemic as mutations continue to accumulate^12,13^. Continued identification and evaluation of mutants is crucial in order to better understand the evolving nature of the pandemic, and to ensure that the treatments and vaccines whose primary target is the Spike protein continue to be effective. As part of a large collaboration to review and characterize the evolution of the SARS-CoV-2 proteome in three-dimensions, an analysis of the SARS-CoV-2 genomes deposited into the GISAID database^14^ at the end of June 2020 was conducted. A full description of the methods used to analyze the mutations of all of the SARS-CoV-2 proteins, including the Spike protein, can be found elsewhere^15^, and the raw data is made freely available^16^. Based on that analysis of 33,290 viral genomes, there are several interesting trends in the mutations accumulated in the Spike protein RBD. First, 444 (1.3%) contained a mutation in the RBD of the Spike protein; of these, 144 unique sequence variants were identified. The identified mutations account for substitutions of 78 individual residues in the RBD (residues: 330-527), with the top 5 substitutions listed in **Fig. 2c**. Among the flexible loop regions, Loop 1 (residues 438-450) contains 5 unique mutations, Loop 2 (residues 455-470) contains 9 unique mutations, Loop 3 (residues 471-491) contains 21 unique mutations, and Loop 4 contains 3 unique mutations. While all of the flexible regions of the RBD have residues that have been found to be mutated in the current COVID-19 pandemic, Loop 3 seems to be a particular hotspot of mutation, with 13 out of 20 residues having at least 1 mutation identified. The top 4 most common mutants of Loop 3, based on number of genomes containing these mutants, were chosen to be studied in more detail: T478I, S477N, V483A, G476S (**Fig. 2d**).

### MD Simulations Reveal that the Flexible Loop 3 of CoV2-RBD is Resilient to Localized Mutations

Based on the 4 common mutations identified in Loop 3 of the RBD, MD simulations of the single mutants G476S, S477N, T478I, or V483A were performed to observe how they affect the RBD’s conformation and dynamics. Using the same starting crystal structure as the wildtype simulation (PDB: 6m0j), 4 new starting structures were created by mutating the relevant residue in pyRosetta^17^. These new structures were then subjected to the same energy minimization and equilibration conditions as the wild-type structure (see Methods), before collecting 2 μs-long MD simulations for each under the same conditions as used for the wildtype simulation. Analysis of the backbone RMSD over the time-course of each simulation (**Fig. S2b**) shows that all of the mutant structures remain in a relatively stable equilibrium from their respective starting points (average RMSD: G476S 1.65 Å; S477N 1.48 Å; T478I 1.46 Å; V483A 1.44 Å). A closer look at the fluctuations of the backbone atoms again illustrates similar conformations as observed for the wild-type simulation through the MD snapshots (**Fig. 5a**). The per-residue RMSF profiles of the mutant simulations show that there is no significant difference in backbone flexibility between the 4 mutants in Loop 3 (**Fig. 5b**), although T478I appears to be marginally more perturbative than the other mutants, slightly increasing the flexibility of the Loop 3 region. The average structures show that there is virtually no difference between the backbones of the wild-type or mutants (**Fig. 5c**). This suggests that the conformational flexibility of Loop 3 is resilient to single mutations, and this resiliency may account for the higher number of mutations observed in this region.

**Figure 5.**
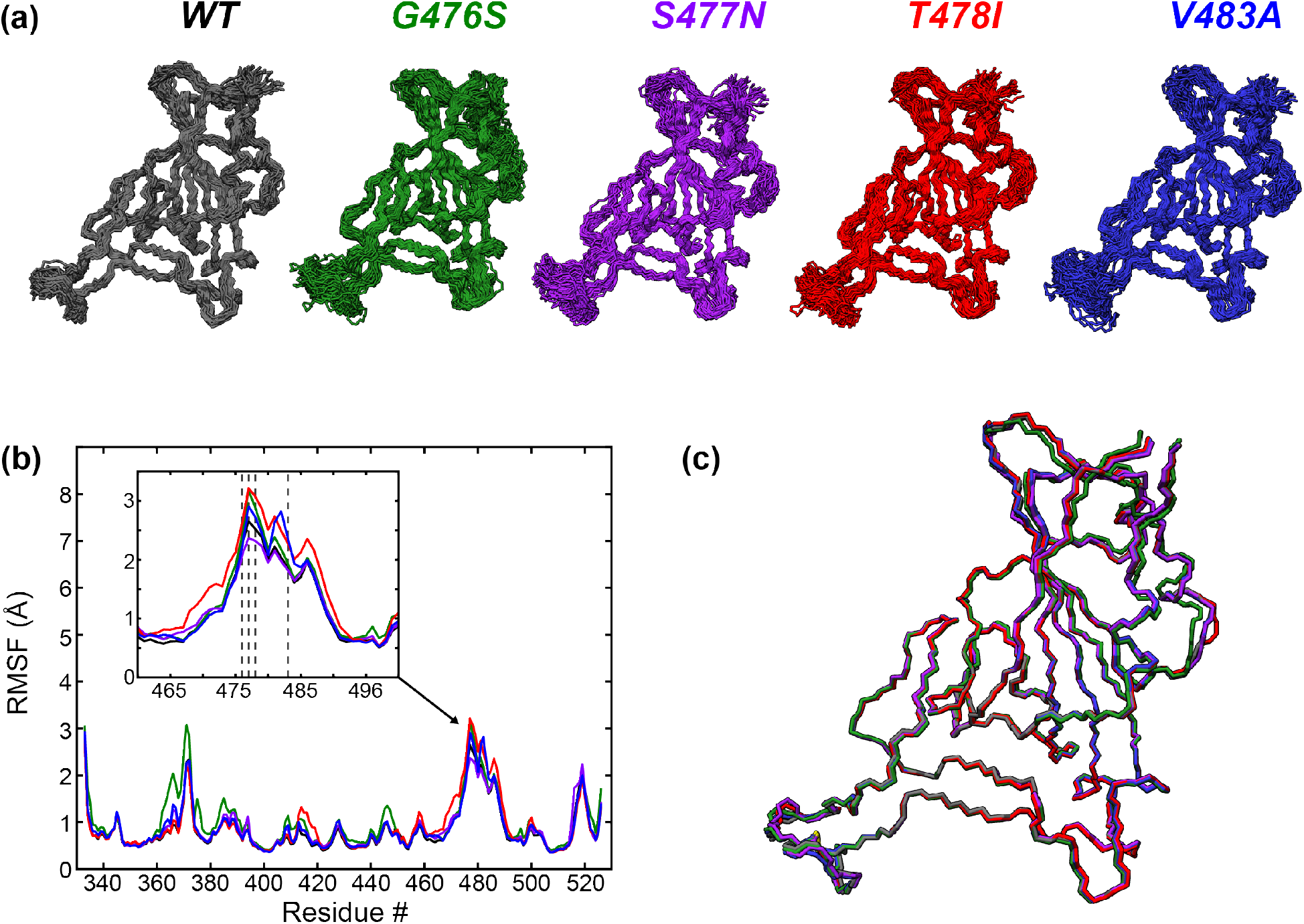
MD simulations of 4 mutants in the flexible Loop 3 region of the SARS-CoV2 RBD binding interface. (a) Snapshots of conformations sampled during 2 μs MD simulations of mutants G476S (green), S477N (purple), T478I (red) and V483A (blue). The wild-type snapshots (black) are the same as shown in Figure 3b, reproduced here for comparison with the mutants. (b) Per residue RMSF of all backbone (N, CA, C) atoms from the 5 different models. The inset shows the Loop and Loop 3 regions. RBD mutations are clustered in this region (dashed lines: G476S, S477N, T478I, V483A). (c) Average conformations of the 5 different models, showing the high similarity between all of the models. The colors of the data and models are kept consistent throughout the figure.

### Cluster Analysis of the RBD Conformational Ensembles from MD Trajectories

To better examine the conformational states of the RBD binding interface that were sampled during the MD simulations, and to identify binding and non-binding conformational states, we performed a cluster analysis on each of the wild-type and mutant simulations using a hierarchical agglomerative (heiragglo) algorithm^18^. Using an epsilon (e) cutoff of 1.9 Å, the 22500 conformations of the CoV1-RBD and CoV2-RBD were separated into 51 and 14 clusters respectively, whereas the CoV2-RBD mutants clustered into fewer? groups (G476S: 4 clusters, S477N: 4 clusters, T478I: 2 clusters, V483A: 7 clusters). The average RMSD of the residues in the RBD binding interface with respect to the starting crystal structure was then calculated for each cluster. Clusters with low RMSD then represent conformations that are very similar to the crystal structures of RBD bound to ACE2, while clusters with large RMSD correspond to conformations that are very different from these receptor bound states. **Figure 6** shows the RBD binding interface of the average conformation of each cluster with the smallest RMSD (ie. most similar to the bound state) in blue and the largest RMSD (ie. least similar to the bound state) in pink for each of the MD simulations presented. Interestingly, the biggest difference in conformation is observed with the structures of the largest RMSD clusters of the loop-models from both CoV1-RBD and CoV2-RBD, where a large portion of Loop 3 is curled back over the binding interface (**Fig. 6a,b**). This conformation of the free RBD may block the binding interface and prevent interactions with ACE2.

**Figure 6.**
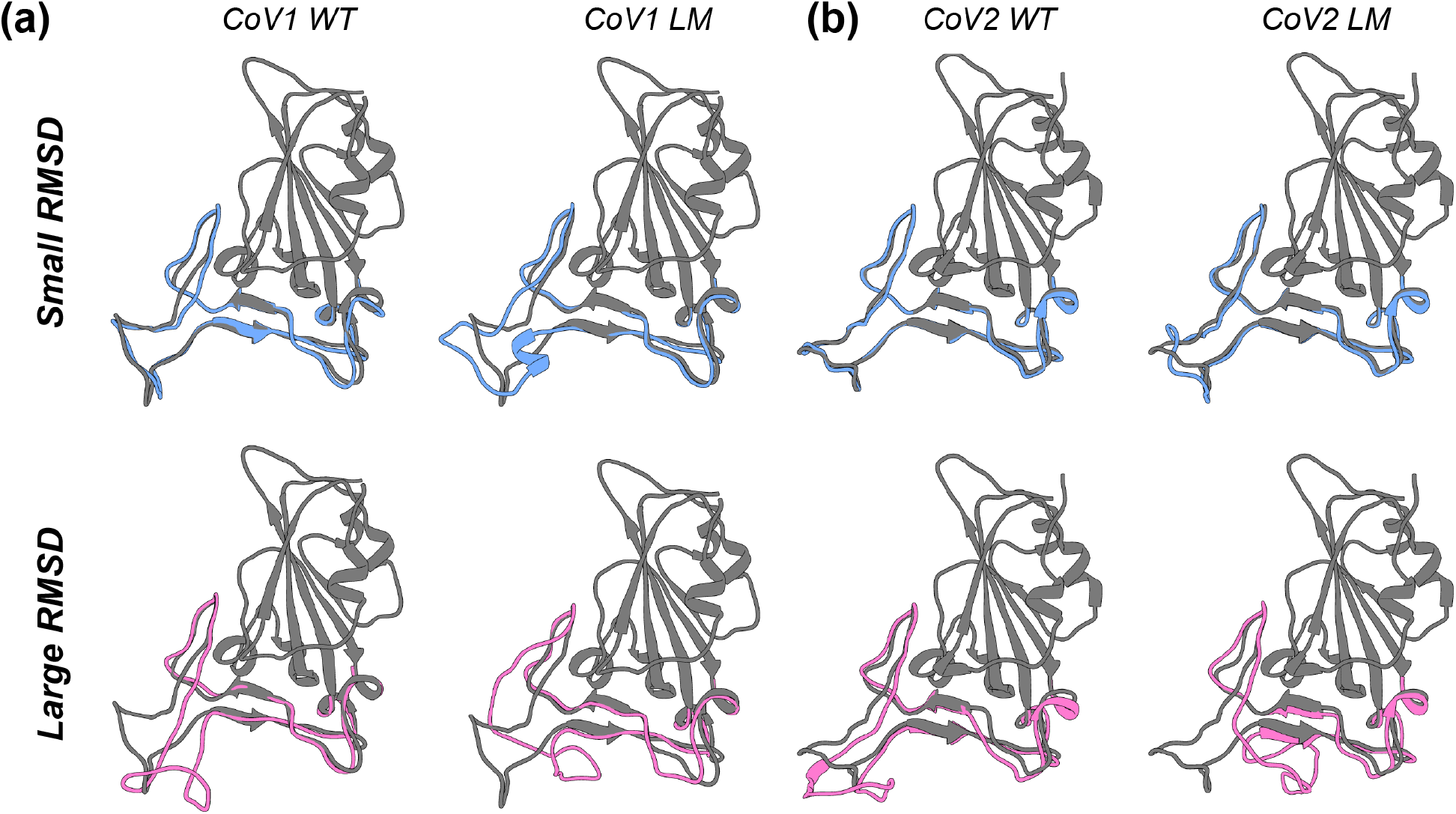
Representative conformations from MD simulations that show the greatest similarity or difference from the starting crystal structure of CoV1-RBD or CoV2-RBD. (a) The conformations that are most similar (blue, small RMSD) or most different (pink, large RMSD) of the binding interface loops from the wild-type (left) or loop-modeled (right) CoV1-RBD. The crystal structure (PDB: 2dd8) is shown in dark gray. (b) The conformations that are most similar (blue, small RMSD) or most different (pink, large RMSD) of the binding interface loops from the wild-type (left) or loop-modeled (right) CoV2-RBD. The crystal structure (PDB: 6m0j) is shown in dark gray.

## 3. Discussion

Because of the importance of the spike RBD in the initial binding of the SARS-CoV2 viral particle to a host cell, it is important to have an understanding of the conformation and dynamics at all stages of the binding event, including in the unbound state. In particular, modeling and identifying conformations of the free state are informative on a range of conformations that can be targeted by therapeutics. However, many of the RBD structures that have been determined and deposited into the PDB are either incomplete, mainly missing residues in the loop sections of the RBD, or are in a bound state in complex with the ACE2 receptor or various neutralizing antibodies. Much of the large binding interface of the RBD does not adopt strong secondary structure elements, but are rather random coil loops. These loop regions (**Fig. 2a**) are not predicted to fall under the definition of intrinsically disordered regions (**Fig. 2b**). However, it is interesting to note that the loops tend to have higher predicted disorder propensity than the rest of the RBD (**Fig. 2b**), a propensity that may be evolutionarily conserved^19^. In addition, the missing residues of these loop regions in many cryo-EM structures of the spike protein suggest that these loops may be dynamic and sample a conformational ensemble distinct from the bound state. Indeed, in all of the structures investigated in this work the residues within the loop regions show large RMSF values, with Loop 3 having the largest. The fact that Loop 3 is so flexible is quite interesting since this loop is directly adjacent to the binding interface with ACE2, and even provides some stabilizing contacts to residues on ACE2. In order to probe the conformational flexibility of Loop 3 and characterize the unbound conformational space of the RBD beyond the ACE2 bound state, it was necessary to perturb the loop away from the stable low-energy state of the RBD-ACE2 crystal structure. These loop models again showed very high flexibility of Loop 3 (ie. large RMSF), while maintaining the same average flexibility in other regions of the RBD as observed in the simulation starting from the CoV1 or CoV2 crystal structures (compare **Fig. S3** with **Fig. 3a**). This indicates that the bulk of the RBD structure is resilient to change even in the presence of large conformational flexibility of Loop 3. In addition, some of these loop models represent conformations that are more energetically favorable for the unbound RBD in solution (**Table 1**) on average.

The large and relatively flat binding interface between the RBD and ACE2 represents an interesting protein-protein interaction that provides a challenging target for traditional small molecule therapeutics that typically bind to well-defined binding pockets on targets such as enzymes. Instead, with the conformational flexibility afforded to the binding interface of the RBD the identification of lowly populated or transient cryptic binding sites should be considered. Cryptic binding sites are difficult to determine in the unbound apo state of a protein, but are generally found in and around dynamic and flexible protein regions, where the inherent conformational fluctuation allows for cryptic sites to become accessible^20–23^. By comparing the conformations that the CoV1-RBD and CoV2-RBD sampled during our MD simulations to the corresponding crystal structure of ACE2 bound RBD, we were able to identify conformations of the dynamic and flexible loop regions that were distinctly different from the bound state of the RBD. In particular, the MD simulations of the different loop models sampled conformations of the CoV2-RBD that contained stabilizing interactions between the sidechain of Q493 and the backbone of F486, helping to fold Loop 3 over the binding interface of the RBD (**Fig. 6b**) and which would block the normal RBD binding interface with ACE2. Such examples of conformations that can be sampled by the RBD ensemble in solution, which provide natural interruption of the protein-protein binding interface between RBD and ACE2, represent potential targets that create transient and/or cryptic binding sites that can be exploited by therapeutic design.

The impact of mutations on the structure and conformational flexibility of the spike protein, especially during an ongoing pandemic, is of particular concern when designing therapeutics against SARS-CoV-2. For example, based on recent structures of the D614G spike protein obtained by cryo-EM it is now becoming clear that the D614G mutation interferes with a stabilizing interaction between monomers of the trimeric assembly^24^, providing increased infectivity of the virus by ensuring that all three of the RBDs in the spike protein have the flexibility to adopt binding-competent, open conformations^24–26^. There is thus a persistent need to maintain a current understanding of the impact of mutations that are manifesting during the course of the current pandemic. Several common RBD mutants have been identified previously and their effects on binding to ACE2 have been probed, including N439K, T478I, V483A, G476S, S494P, V483F, and A475V^12^. Ghorbani et al. characterized these mutants in the context of the full RBD-ACE2 complex (PDB: 6m0j) through molecular dynamics simulations, showing a stable and overall similar RMSD among the wild-type and mutants in the extended loop forming the binding interface with ACE2^12^. The binding free energy between the RBD and ACE2 was also found to be consistent between the wild-type and the mutants with the exception of T478I, which had a binding free energy 6 kcal/mol higher than the wild-type. These results are consistent with experimental data from deep mutational scanning and flow cytometry, which found that all naturally occurring mutants have a similar degree of expression and a similar binding affinity for ACE2 as in the wild-type^13^. Our own analysis of the SARS-CoV-2 proteome evolution^15^ during the current pandemic has identified G476S, S477N, T478I, and V483A as mutations that have appeared in the RBD and are clustered within Loop 3 (**Fig. 2d**). Similar to the simulations of the RBD-ACE2 complex^12^, the MD simulations of the mutants of the unbound RBD in this work do not show large perturbing effects on the average conformational state. This suggests that while these mutants may have an impact on the stability of the binding interface with ACE2, they do not greatly perturb the conformational state of the RBD in solution and may even serve to reduce the conformational sampling of the unbound RBD.

The ongoing COVID-19 pandemic caused by SARS-CoV-2 has focused the collective scientific community to quickly provide both knowledge and action to help alleviate the effects of this crisis. In this work, our data indicate that common mutations identified in the Loop 3 region of the CoV2-RBD are fairly non-perturbing and do not affect its conformational flexibility and sampling in an unbound solution state, suggesting a therapeutic designed to target this region may be broadly applicable to RBDs with mutations in this region. In addition, we have identified unique conformations of the unbound CoV2-RBD in solution that naturally block the binding interface with ACE2 and may be interesting targets for drug design to interfere with RBD-ACE2 binding. We hope that these results will help to catalyze future identification of therapies relevant to CoV2 or to future coronaviruses that may emerge.

## 4. Materials and Methods

### Preparation of Initial Structural Models

The structures used to model the wild-type RBD’s were taken from the Coronavirus Structural Taskforce (https://github.com/thorn-lab/coronavirus_structural_task_force), which further refined the high-resolution structures determined by X-ray diffraction of CoV1-RBD in complex with a neutralizing antibody (PDB: 2dd8) and of CoV2-RBD in complex with the human ACE2 receptor (PDB: 6m0j). In order to isolate the RBD for subsequent MD simulations, the protein modelling platform Pyrosetta^17^ was employed to remove the ACE2 receptor residues and RBD glycans from the model, leaving only the clean RBD residues. The 4 selected mutants (G476S, S477N, T478I, V483A) were then generated from the clean wild-type RBD structure by creating a decoy of the wild-type structure in Pyrosetta and restricting for the selected mutation. These mutant decoys were then relaxed based on the ref2015_cst score function within Pyrosetta^27^. One-hundred energy minimized decoys for each mutant were generated in this protocol, and the lowest energy decoy for each mutant RBD was selected as the starting structure for MD simulation.

### Rosetta Loop Modelling

Loop 3 variant structures of the wild-type RBD were generated using the lowest energy decoy of the wild-type RBD, using the same protocol as described for the mutant models. The loop being modelled was defined from residues 472-490 in Pyrosetta with jumps in the foldtree introduced at residue 470, 481, and 492. The Pyrosetta KinematicMover^17,27^ was then used to search for a different conformation in the loop carbon backbone with residues 472 and 490 as pivots. Only conformations maintaining the critical disulfide between C480 and C488 were selected to output a decoy, and this protocol was run until 100 decoys had been generated. All 100 decoys were then relaxed based on the ref2015_cst score function using Pyrosetta^27^. Once again, 100 energy minimized structures for each initial loop decoy were generated and 5 loop structures were chosen at random for Molecular Dynamics simulations. The lowest energy decoy of each of the 5 loop structures was used for MD simulation.

### Molecular Dynamics Simulations

All of the water molecules in the initial X-ray structure were removed. Each protein was immersed in a truncated octahedral box of OPC water molecules^28^ with the box border at least 20 Å away from any atoms of the RBD. Each system was neutralized by adding counter ions (Cl^-^ or Na^+^). The protein was treated with the ff19SB force field^29^. The simulations were performed with the GPU-enabled CUDA version of the pmemd module in the AMBER 2018 package^30^. Prior to MD simulation, the systems were subjected to energy minimizations and equilibration. The minimization started with 1000 steps of steepest descent minimization followed by 1000 steps of conjugate gradient minimization. The system was heated from 0 to 300 K over 100 ps with protein position restraints of 10 kcal mol^-1^ A^-2^. Then a series of equilibrations (each lasting 10 ns) were performed at constant temperature of 300 K and pressure of 1 atm with protein position restraints that were incrementally released (10.0, 1, 0.1 and 0 kcal mol^-1^ A^-2^). Periodic boundary conditions were used, and electrostatic interactions were calculated by the particle mesh Ewald method^31,32^, with the non-bonded cutoff set to 9 Å. The SHAKE algorithm^33^ was applied to bonds involving hydrogen, and a 2 fs integration step was used. Pressure was held constant at 1 atm with a relaxation time of 2.0 ps. The temperature was held at 300 K with Langevin dynamics and a collision frequency of 5.0 ps^-1^. The production runs for wild-type CoV1-RBD and wild-type CoV2-RBD are 4 μs, for the CoV2-RBD mutants are 2 μs, and for the CoV1-RBD and CoV2-RBD loop models are 750 ns.

All analysis of MD trajectories, including the root-mean-square deviation (RMSD), rootmean-square fluctuation (RMSF), hierarchical agglomerative clustering, Molecular Mechanics/Generalized Born Surface Area (MM/GBSA) free energy calculation, and the extraction of representative structures from trajectories were performed using CPPTRAJ^34^ as implemented in AMBER18. Visualization of structures was performed with UCSF Chimera^35^. Cluster analysis was performed on the binding interface towards ACE2 of SARS-CoV-RBD and SARS-CoV2-RBD (residues 432-492 for SARS-CoV-RBD and residues 445-506 for SARS-CoV2-RBD, respectively) using the average-linkage hierarchical agglomerative method^18^. Coordinate RMSD was used as the distance metric. The critical distance ε value was set to 1.9 Å and the sieve value was set to 10. Only the backbone C, CA, and N atoms were used in the clustering.

## Data Availability

All data and protocols are available upon reasonable request to the corresponding author. All data used for mutational analysis is freely available in^15^ and^16^.

## Author Contributions

JKW, BW, AS, DAC and JB designed research. JKW, BW, and AS performed research. JKW, BW, AS, and CLH analyzed data. All authors contributed to writing and editing the manuscript.

## Acknowledgements

This work was supported by a National Institutes of Health Grant GM136431 (JB) and Rutgers University Center for COVID-19 Response and Pandemic Preparedness (CCRP2) research grants (JB, DAC).

