## Supporting Information for "Molecular Dynamics Analysis of a Flexible Loop at the Binding Interface of the SARS-CoV-2 Spike Protein Receptor-Binding Domain"

**Contents:**
Figure S1-S3

(a) Loop 3 Contacts (b) No Loop 3 Contacts

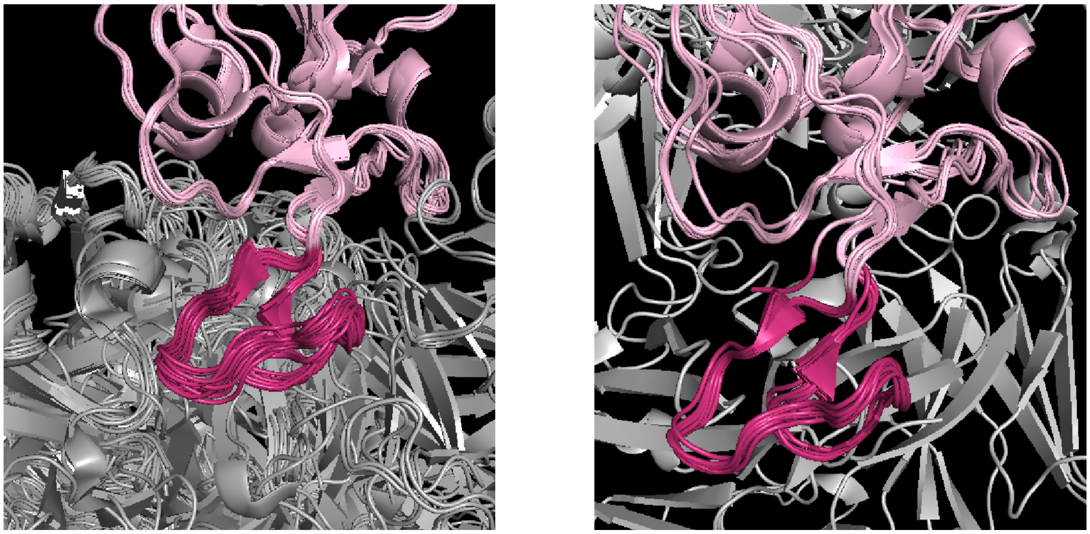

**Figure S1. Antibody-Bound Structures of the CoV2-RBD in the PDB.** (a) Overlay of structures deposited into the PDB with antibodies that contact Loop 3 of the CoV2-RBD (PDB: 6xc2, 6xc3, 6xc4, 6xc7, 6xe1, 6xkq, 7bz5, 7cdi, 7cdj, 7ch4, 7ch5, 7chb, 7che, 7chf, 7cjf, 7jmo, 7k9z, 7k45). (b) Overlay of structures deposited into the PDB with antibodies that bind to the RBD at locations other than Loop 3 (PDB: 6w41, 6xkp, 6yla, 6ym0, 6zdg, 6zer, 6zfo, 7cah, 7jmw, 7jva, 7jx3). In both panels, structures were aligned to the RBD domain of the CoV2-RBD bound to ACE2 (PDB: 6m0j).

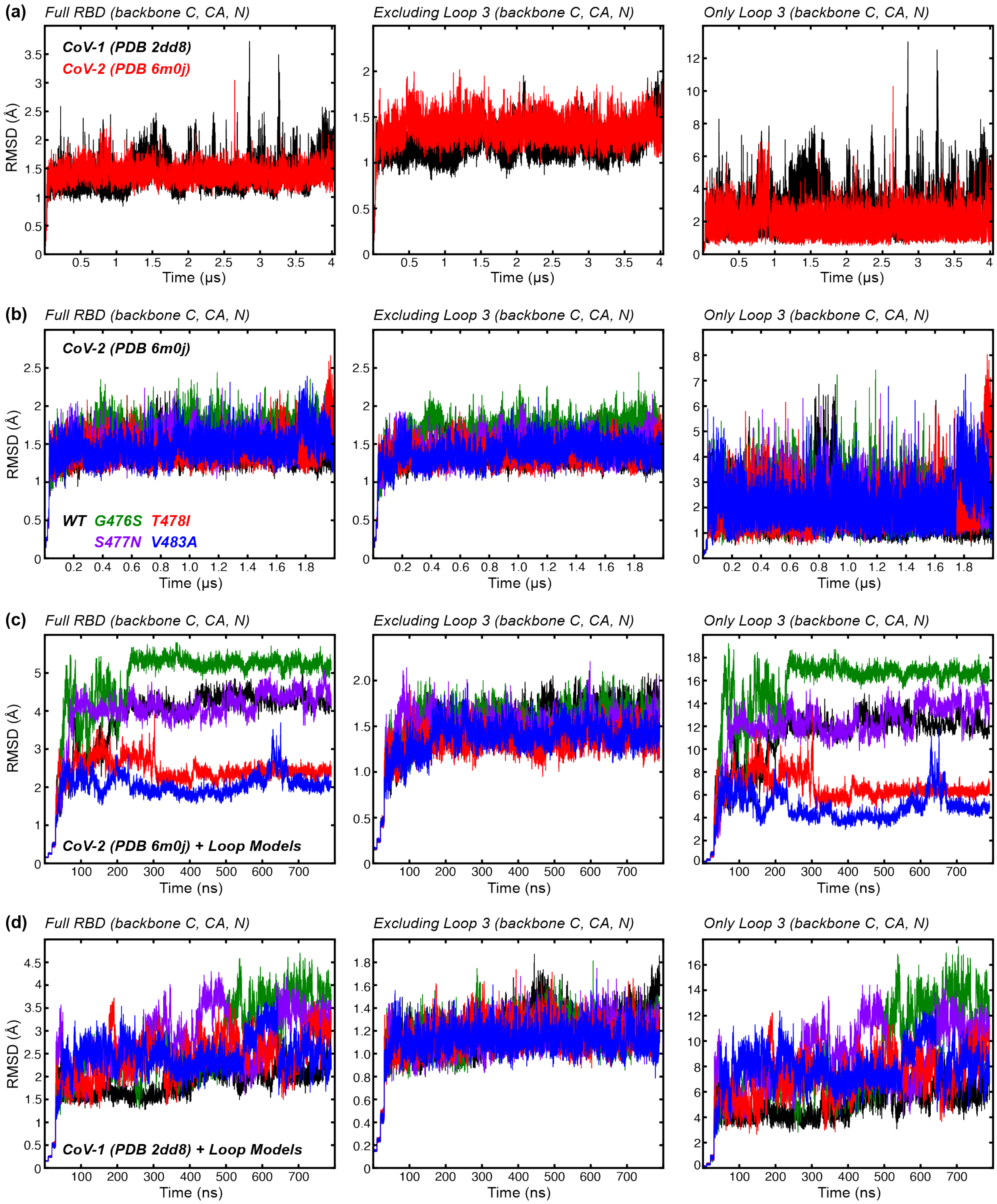
(a)
Full RBD (backbone C, CA, N)
Excluding Loop 3 (backbone C, CA, N)
Only Loop 3 (backbone C, CA, N)
CoV-1 (PDB 2dd8)
CoV-2 (PDB 6m0j)
RMSD (Å)
Time (μs)
(b)
Full RBD (backbone C, CA, N)
Excluding Loop 3 (backbone C, CA, N)
Only Loop 3 (backbone C, CA, N)
CoV-2 (PDB 6m0j)
WT G476S T478I
S477N V483A
RMSD (Å)
Time (μs)
(c)
Full RBD (backbone C, CA, N)
Excluding Loop 3 (backbone C, CA, N)
Only Loop 3 (backbone C, CA, N)
CoV-2 (PDB 6m0j) + Loop Models
RMSD (Å)
Time (ns)
(d)
Full RBD (backbone C, CA, N)
Excluding Loop 3 (backbone C, CA, N)
Only Loop 3 (backbone C, CA, N)
CoV-1 (PDB 2dd8) + Loop Models
RMSD (Å)
Time (ns)

**Figure S2. Root-mean-square deviation (RMSD) of the backbone (N, CA, C) atoms relative to the starting structures. The left column is the RMSD of the full RBD, the center column is the RMSD excluding the residues of Loop 3, and the right column in the RMDS considering only the residues of Loop 3.** RMSD of conformations sampled from: (a) MD simulations of SARS-CoV RBD (black) and SARS-CoV2 RBD (red); from (b) MD simulations of CoV2-RBD mutants G476S (green), S477N (purple), T478I (red), and V483A (blue); from (c) MD simulations of different loop models of CoV2-RBD; and from (d) MD simulations of different loop models of CoV-RBD. The colors used in the RMSD plots match to the same colored structures and RMSF plots of the main text.

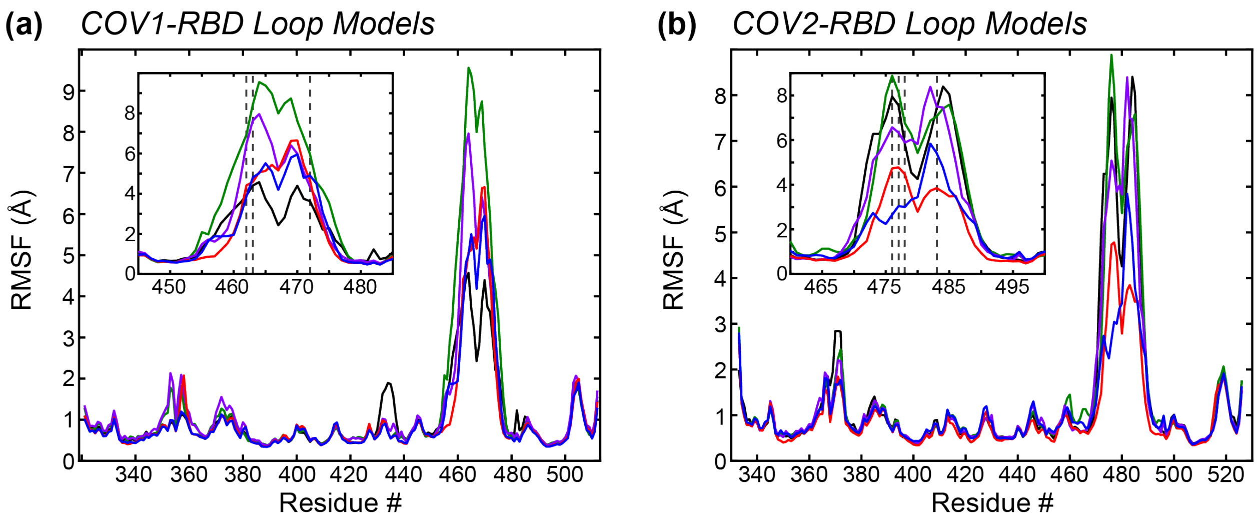


**Figure S3. Per-residue root-mean-square fluctuations (RMSF) of the backbone C, CA, and N of different loop models.** (a) RMSF plots from the 5 different CoV-RBD loop models. (b) RMSF plots from the 5 different CoV2-RBD loop models. The colors used here match with the models used in Figure 5 of the main text.
